# Caudate nucleus aging and its association with brain disorders

**DOI:** 10.64898/2026.10.28.762543

**Authors:** Shuai Guo, Xingyi Yang, Yi Dong, Jixiang Xu, Angela Sangjizhuoma, Thomas Wichmann, Adriana Galvan, Yoland Smith, Jing He

## Abstract

The caudate nucleus supports cognition and goal-directed behavior, undergoes age-associated changes, and is implicated in substance use, psychiatric, and neurodegenerative disorders. However, how aging alters its diverse cell types and how such changes relate to diseases remains poorly understood. To address these questions, we assembled single nucleus transcriptomic atlases of the caudate tissue, comprising 963,842 nuclei from 178 human donors, including controls and individuals with alcohol use disorder (AUD) or opioid use disorder (OUD), and 83,699 nuclei from six healthy rhesus macaques. We also curated 3,139 bulk RNA-seq profiles from the caudate and other brain regions spanning controls and multiple diagnostic groups, as well as paired RNA and chromatin accessibility profiles from 257,524 caudate nuclei across 161 control and AUD donors. These datasets enabled comparisons across age, species, brain regions, and diagnostic groups. We found shared glial aging features in humans and macaques, whereas neuronal changes were more divergent. Inferred neuron-to-neuron communication declined with age, with greater declines in AUD. Both transcriptional and chromatin accessibility profiles showed nonlinear age-associated patterns, with pronounced changes in later adulthood. In donors with AUD, age-associated changes appeared earlier, with altered trajectories in lipid metabolism, mitochondrial function, and neuronal signaling. Across comparisons involving AUD, OUD, bipolar disorder, schizophrenia, and Alzheimer’s disease, mitochondrial function and fatty acid metabolism emerged as consensus features linking caudate aging to diseases, although the direction and extent of overlap varied by disorder and brain region. Together, these findings define cellular and regulatory features of primate caudate aging and reveal shared and context-dependent relationships between aging and brain disorders.

## Introduction

Aging involves the accumulation of cellular damage that contributes to functional decline and increased vulnerability to disease^1^. In the brain, aging is accompanied by region- and cell type-specific alterations in cellular composition, gene expression, and intercellular communication^2^. Much cell type-resolved research on brain aging has focused on the cortex^3–6^, leaving subcortical regions less well characterized. The caudate nucleus, the associative division of the dorsal striatum, supports cognition^7^ and goal-directed actions^8,9^. Most of its neurons are GABAergic medium spiny neurons (MSNs)^10–12^, broadly organized into *DRD1*-expressing direct pathway and *DRD2*-expressing indirect pathway populations^13^ within striosome and matrix compartments^14^. With age, the caudate nucleus volume decreases^15^, and the availability of dopamine receptors and dopamine transporters declines^16,17^, whereas the expression of complement and other neuroinflammatory genes increases^18^. How these changes are distributed across caudate nucleus cell types, how they change with age, and whether they are conserved across primates remains poorly understood.

Understanding caudate nucleus aging is also relevant to disorders involving striatal circuitry. Early striatal amyloid deposition occurs in presenilin-1 mutation carriers, i.e., individuals at risk of developing early Alzheimer’s disease (AD), and the caudate nucleus volume is reduced in cases of AD, regardless of etiology^19,20^. Striatal dopamine synthesis capacity is elevated in schizophrenia (SCZ) and bipolar disorder (BIP) with psychosis^21–23^. Substance use disorders also involve altered striatal cue processing and dopamine signaling, including greater dorsal striatal cue reactivity in those with heavy alcohol use, and reduced stimulant-evoked striatal dopamine release in individuals with alcohol or heroin dependence^24–29^. Aging may intersect with these disease-associated changes. MRI-estimated brain-age gaps are larger in dementia, SCZ, and BIP^30^, as well as alcohol use disorder (AUD)^31^, in which the gap increases in later decades^32^. Striatal disease signatures also share immune and mitochondrial features with aging^33,34^. These observations raise the question of whether disease-associated molecular changes follow, oppose, or modify normal aging trajectories.

Single nucleus studies have begun to characterize these relationships at the cellular level. A previous caudate nucleus study linked a higher proportion of inflammatory microglia to both AUD and older age^28^, while a separate study identified opioid use disorder (OUD)-associated changes in neuronal inflammatory and DNA damage-related pathways^29^, which are canonical hallmarks of aging^35^. However, it remains unclear whether these disease-associated changes align with normal aging or reflect altered aging trajectories. Addressing this question requires cell type-resolved references for normal aging and explicit tests of whether age associations differ by diagnosis. A related question is whether transcriptional changes are accompanied by changes in chromatin accessibility. Previous studies reported relatively limited age-associated accessibility changes within human midbrain cell types^36^, whereas regions that became more accessible with age in mouse cell types were enriched for activator protein 1 (AP-1) motifs^37^. How transcriptional and chromatin changes relate to one another during caudate aging, and how disease modifies these relationships, remains unresolved.

Here, we combined human and macaque caudate single nucleus RNA sequencing (snRNA-seq) atlases with bulk transcriptomic and paired chromatin accessibility data to characterize aging and its relationship to species, brain regions, and brain disorders. We identified species-independent (shared) glial aging features, along with species-dependent neuronal changes, and defined nonlinear molecular trajectories across human adulthood. We found that AUD was associated with altered timing and direction of these trajectories, accompanied by coordinated changes in chromatin accessibility and activity- and stress-responsive gene expression. Across independent bulk brain cohorts, disease-associated expression changes aligned with, opposed, or showed little correspondence with normal aging, depending on the disorder and brain region. In the caudate samples from individuals with AD, reduced oxidative phosphorylation, respiratory-chain, and MSN-derived calcium/calmodulin signaling programs were also associated with poorer cognitive performance. These findings establish a comparative reference for primate caudate nucleus aging and reveal distinct relationships between aging, disease-associated molecular changes, and cognition.

## Results

### A primate caudate nucleus atlas and reference transcriptional programs

To characterize caudate nucleus cell aging and its relationship to brain disorders, we assembled a snRNA-seq atlas of 963,842 nuclei from 178 human donors: 88 healthy controls, 84 individuals with AUD^28^, and 6 with OUD^29^ (Fig. 1, A and B). Ages were available for 173 donors and ranged from 23 to 82 years, with a median of 56 years; 136 donors were male. We also collected snRNA-seq profiles of 83,699 caudate nuclei from six female rhesus macaque monkeys (see Methods), including three young animals aged 5.0 to 6.5 years and three old animals aged 18.0 to 19.5 years (Fig. 1C). Independent clustering and marker-based annotation identified the same 18 cell populations in both species, which we grouped into 14 major cell types for downstream analyses (Fig. 1, A and C). These included six MSN subtypes (D1-matrix, D1-striosome, D2-matrix, D2-striosome, D1/D2 hybrid cells, and D2 striosome/matrix hybrid), five interneuron subtypes (*PTHLH*+, *SST*+, *CHAT*+, *VIP*+, and *TAC3*+), and seven non-neuronal populations (astrocytes, oligodendrocytes, oligodendrocyte precursor cells (OPCs), microglia, endothelial, mural, and ependymal cells), similar to what has been reported^11,12^. Cell type composition was broadly similar between species (Fig. 1D).

**Fig. 1.**
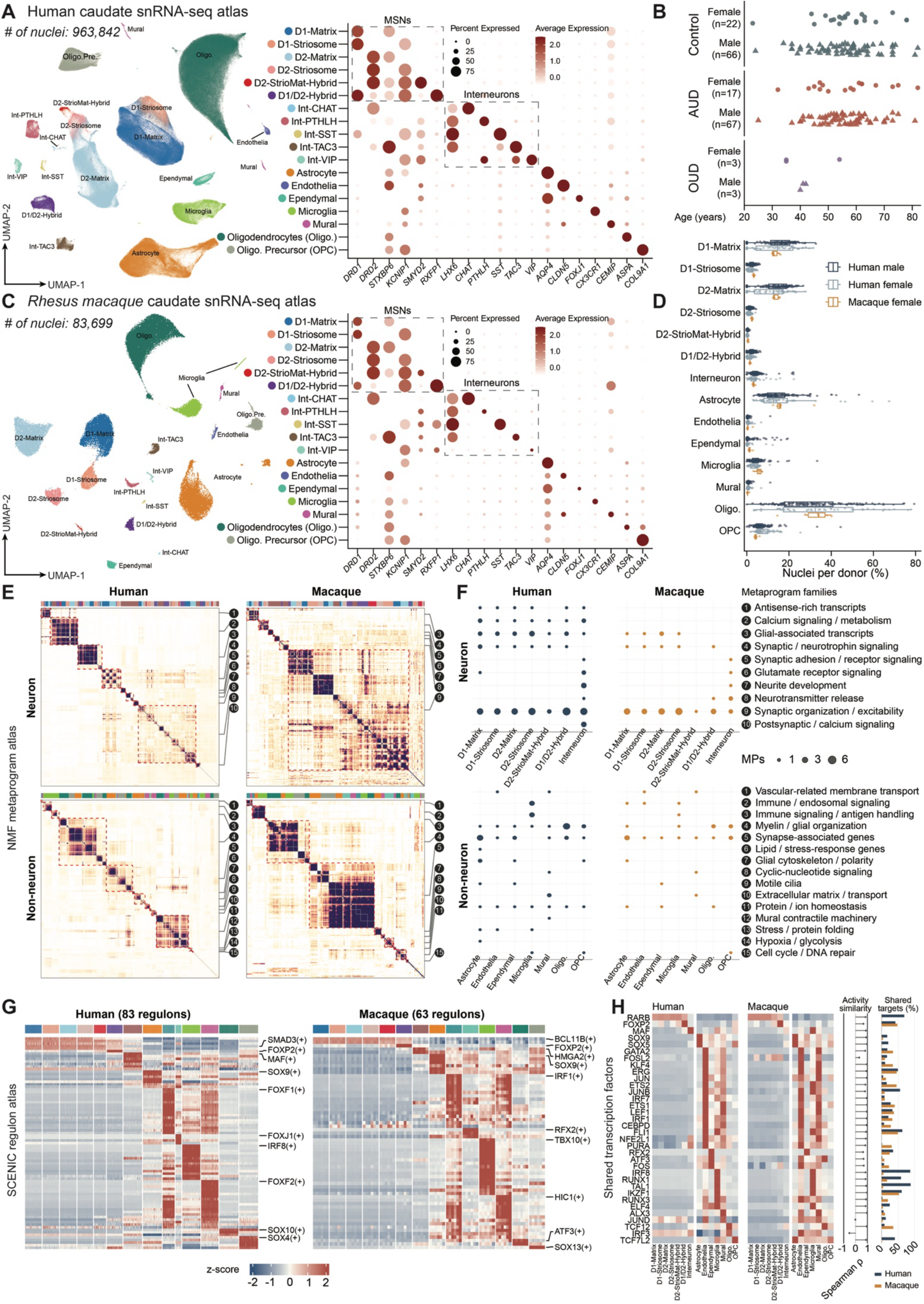
Human and macaque caudate nucleus atlases define reference transcriptional programs. **(A)** Human caudate snRNA-seq atlas and marker expression. UMAP colors indicate cell populations. In the dot plot, size indicates the percentage of nuclei expressing each gene, and color indicates average expression. **(B)** Human donor ages by diagnosis and sex. Circles represent females and triangles represent males. Ages are available for 173 donors; three controls and two AUD donors lack recorded ages. **(C)** Macaque caudate atlas from three young and three old females, displayed as in **(A)**. **(D)** Cell type proportions among recovered nuclei per donor, shown for human males (*n* = 136), human females (*n* = 42) and macaque females (*n* = 6). Points represent donors; boxes show medians and interquartile ranges. **(E)** Pairwise Jaccard similarity of NMF programs, each represented by its 50 highest-loading genes. Gray and dashed red outlines mark MPs and MP families, respectively. Top bars indicate cell type; family numbers correspond to **(F)**. **(F)** Distribution of 131 human and 68 macaque MPs across families and cell types. Dot size indicates the number of MPs. **(G)** Activity of 83 human and 63 macaque regulons across donor by cell type profiles. Colors show within-regulon z-scores of donor-mean AUCell activity; “(+)” denotes an inferred activating regulon. **(H)** Comparison of 34 shared regulons. Heatmaps show donor-averaged activity across 14 cell types, standardized within each regulon and species. Adjacent plots show cross-species Spearman correlations of activity profiles and shared targets as a percentage of each species’ target set.

To identify expression programs within each cell type, we applied consensus non-negative matrix factorization (cNMF) to each cell type in each individual. This analysis identified 10,550 robust programs in humans and 378 in macaques. Within each species and cell type, we grouped similar robust programs observed across donors into metaprograms (MPs)^38,39^ (see Methods). This yielded 131 human and 68 macaque MPs. Each MP was defined by 50 genes most frequently found among its element robust programs. We then clustered MPs by gene-set overlap, separately for neuronal and non-neuronal cells, to organize related programs across cell types and species. We identified10 neuronal and 15 non-neuronal MP families (Fig. 1, E and F).

Synaptic organization and excitability constituted the largest family of neuronal MPs in both species, comprising 30 human and 22 macaque MPs and including genes such as *DLGAP2*, *GNG7*, and *CACNA1C*^40^ (Fig. 1, E and F). Other neuronal gene families encompassed synaptic and neurotrophin signaling, glutamate receptor signaling, neurotransmitter release, neurite development, and calcium signaling and metabolism. Non-neuronal gene families included vascular-related membrane transport, immune and endosomal signaling, myelin and glial organization, extracellular matrix formation, and cilia motility. Representative genes included *ABCB1* and *SLC7A5* in endothelial and mural cell programs^41^, *MBP* and *ST18* in oligodendrocyte-lineage programs^42^, and *DNAH6* and *DNAH11* in ependymal programs^43^. Seven neuronal and 11 non-neuronal gene families contained MPs from both species (Fig. 1F). Within corresponding cell types, 61 of 131 human MPs (46.6%) and 53 of 68 macaque MPs (77.9%) shared at least 10 of their 50 genes with an MP from the other species, suggesting cross-species consistency.

We next used pySCENIC^44–46^ to infer transcription factor-target networks in caudate cell types (Fig. 1, G and H). Donor-level co-expression networks were aggregated within each species and pruned to motif-supported transcription factor-target relationships, yielding atlas catalogs of 83 human and 63 macaque regulons. We used AUCell^45^ to score regulon activity in each nucleus and identified the regulon characteristics of each cell type (Fig. 1G). In humans, prominent cell type-specific regulons included SOX9 in astrocytes, SOX10 in oligodendrocytes, SOX4 in OPCs, IRF8 in microglia, FOXF1 in endothelial cells, FOXF2 in mural cells, and FOXJ1 in ependymal cells. SOX9 in astrocytes and FOXP2 in D1/D2-Hybrid MSNs ranked highest in their respective cell type in both species^29^. In the other five MSN populations, SMAD3 regulon activity was prominent in humans, whereas BCL11B regulon activity was higher in macaques. We found 34 shared regulons between humans and macaques. After averaging their activities across donors within each cell type, these regulons showed similar activity patterns cross species (i.e., they tended to be more active in similar cell types; median Spearman correlation = 0.87; 31 regulons had correlations above 0.5; Fig. 1H). For 30 of these regulons, the two species shared more predicted target genes than expected under the null model (hypergeometric test, adj. *P* < 0.05). These shared targets accounted for about 16% of the predicted targets per regulon in each species. Thus, the regulators retained some common targets, while most of their predicted targets differ between species. We then built regulon catalogs separately for each cell type, yielding 1,883 human and 1,850 macaque regulon-cell type combinations. Together, the atlas-defined MPs and regulons provide a cell type-resolved reference for transcriptional programs in the primate caudate nucleus.

### Caudate aging shows selective molecular conservation across species

To compare age-associated changes in the human and macaque caudate, we analyzed age-comparable young (n = 6, 23-37 years) and old (n = 13, 66-78 years) human controls alongside three young (5.0-6.5 years) and three old (18.0-19.5 years) macaques (Fig. 2A, see Methods). Cell type proportions were broadly similar between age groups in both species (Fig. 2A). We then compared old and young individuals within each cell type to identify differentially expressed (DE) genes. Non-neuronal cells had more DE genes than neurons (Fig. 2A). Across both neuronal and non-neuronal cell types, 13,161 gene x cell type pairs were significant in at least one species, of which only 577 were significant in both (Fig. 2B). One example shared change was decreased expression of *PELI2* in D2-StrioMat-Hybrid MSNs. *PELI2* encodes an E3 ubiquitin ligase that restrains cGAS-STING signaling, a key regulator in age-associated inflammation and neurodegeneration^47,48^.

**Fig. 2.**
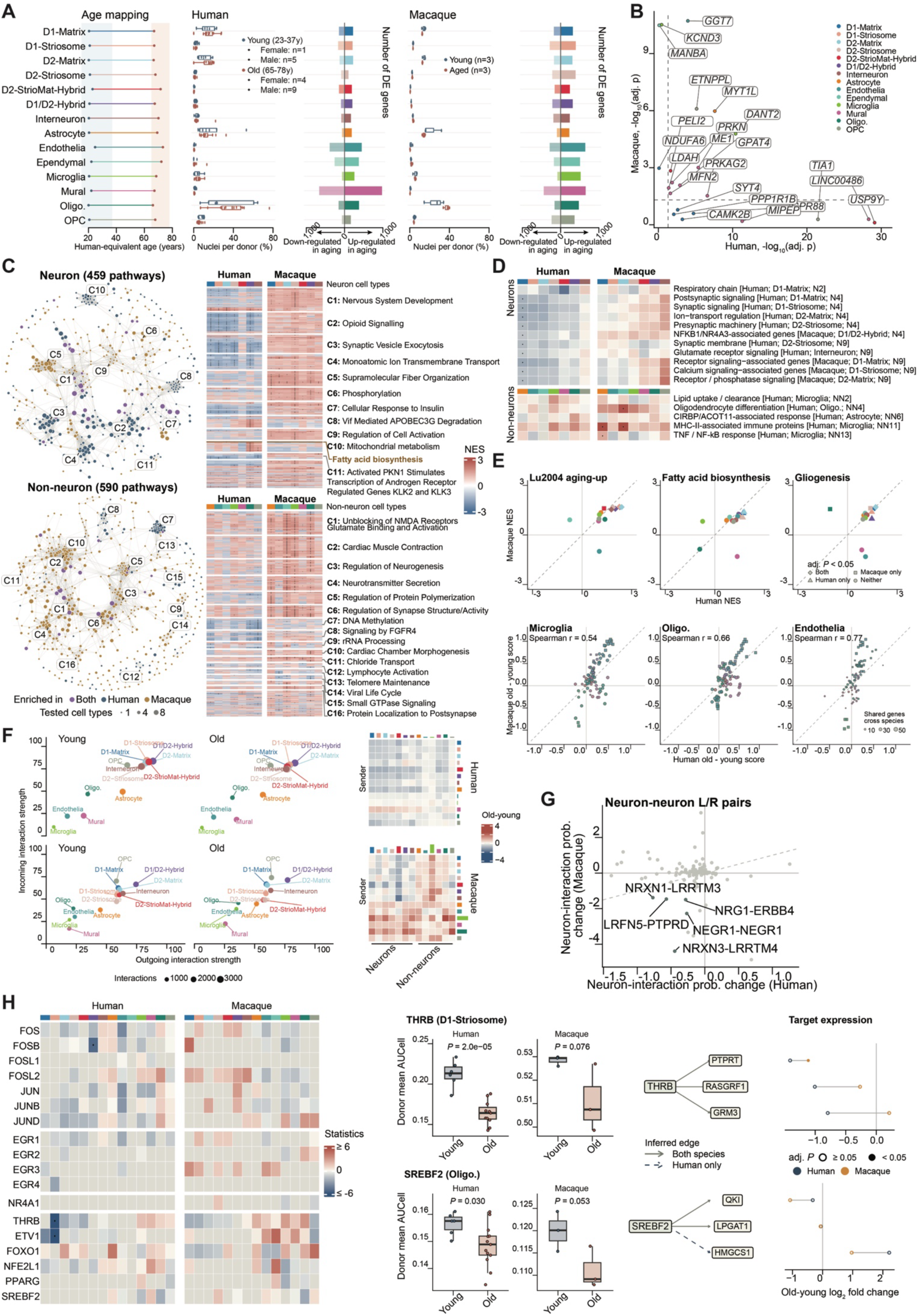
Conserved and divergent features in human and macaque caudate aging. **(A)** Human-equivalent macaque ages, cell type proportions, and DE gene counts. Blue and red indicate young and old groups. Points represent donors; boxes show medians and interquartile ranges; bars show DE genes in old individuals. **(B)** Cross-species comparison of DE genes for gene x cell type pairs. Colors indicate cell type; dashed lines mark adj. *P* 0.05. Selected genes are labeled. **(C)** Pathway enrichment networks and heatmaps. Nodes represent significantly enriched pathways; edges indicate gene-set overlap. Node color indicates species, and size indicates the number of significant species-cell type comparisons. Heatmaps show normalized enrichment scores (NES); dots mark adj. *P* < 0.05. **(D)** Score differences of selected MPs between young and old groups. Colors indicate score differences; dots mark adj. *P* < 0.05, and gray indicates unavailable estimates. **(E)** Cross-species comparisons of selected gene-set NES across 14 cell types (top) and MP score changes in microglia, oligodendrocytes, and endothelial cells (bottom). Colors indicate cell type (top) or MP family (bottom). Bottom point sizes indicate the number of MP genes measured in both species. Shapes indicate significance in each species; dashed lines show equal values. **(F)** CellChat-inferred communication networks. Left, outgoing and incoming interaction strengths; point size indicates interaction count. Right, old-young changes in interaction strength between sender (rows) and receiver (columns) cell types. **(G)** Cross-species comparisons of old-young changes in neuronal ligand-receptor (L/R) interaction probabilities. **(H)** Regulon activity changes with two selected examples. Box plots show donor-mean AUCell activity, alongside selected inferred targets and their old-young expression changes. Heatmap colors show moderated *t* statistics; dots mark adj. *P* < 0.05, and gray indicates unavailable estimates. Solid and dashed edges indicate shared and human-only inferred targets, respectively. Filled points mark DE adj. *P* < 0.05.

We next used gene-set enrichment analysis (GSEA) to compare age-associated pathway changes. The neuronal and non-neuronal pathway networks contained 459 and 590 gene sets, respectively, each enriched in at least one cell type in either species (Fig. 2C). Neuronal enrichment patterns differed substantially between species: synaptic and neurotrophin-related pathways were downregulated in older humans, whereas macaque neurons often showed positive enrichment. Fatty acid biosynthesis and gliogenesis showed positive enrichment trends in both species. Non-neuronal enrichment patterns were more similar between species. Analysis of the atlas-defined MPs showed a similar distinction, with divergent neuronal score changes and more concordant glial changes (Fig. 2D).

To examine whether age-associated changes were shared across species, we compared the enrichment of fatty acid biosynthesis and gliogenesis across cell types in humans and macaques. We also included the canonical aging signature identified by Lu et al.^6^ to assess whether a well-established brain aging signature showed similar patterns across species (Fig. 2E, top). All three gene sets showed broadly similar enrichment patterns across cell types. The lipid-related findings are consistent with reported age-associated changes in lipid composition^49^. At the MP level, old-vs.-young score changes were positively correlated between species in microglia, oligodendrocytes, and endothelial cells, further supporting shared non-neuronal changes (Fig. 2E, bottom).

We then used CellChat to estimate age-associated differences in ligand-receptor communication between annotated cell populations. Neuronal populations generally had higher inferred communication scores than most non-neuronal populations (Fig. 2F). The aggregate neuron-to-neuron score was approximately 5-6% lower in older individuals of both species. Ligand-receptor pairs contributing to shared decreases included NRXN1-LRRTM3 and NRXN3-LRRTM4, which involve synaptic adhesion proteins that organize excitatory synapses^50^ (Fig. 2G).

Age-associated changes in regulon activity were also similar between species for some regulons and different for others (Fig. 2H). THRB regulon activity was lower across the MSN populations in both species. In D1-Striosome MSNs, this decrease was significant in humans and directionally similar in macaques (activity change: human, −0.05, *P* < 0.001; macaque, −0.02, *P* = 0.076). Its inferred targets PTPRT and RASGRF1 also showed lower expression in both species. PTPRT participates in synapse formation, while RASGRF1 contributes to activity-dependent signaling and corticostriatal synaptic plasticity^51,52^. In oligodendrocytes, MP changes were similar across species (Spearman ρ = 0.66; 70 of 85 MPs changed in the same direction). We then looked at the SREBF2 regulon in oligodendrocytes because of its role in regulating cholesterol synthesis and myelin formation^53^. SREBF2 regulon activity decreased in both species (human *P* = 0.030; macaque, *P* = 0.053). *QKI*, which cooperates with SREBF2 during myelin formation^53^, showed lower expression in both species, whereas the cholesterol synthesis gene *HMGCS1*^53^, inferred as a target in humans, showed higher expression in both.

### Human caudate aging follows nonlinear trajectories with late-life changes

To characterize age-associated expression patterns in the human caudate nucleus, we analyzed donor-level pseudobulk profiles from 85 control donors aged 23-82 years. Within each cell type, we calculated Spearman correlations between age and gene expression after adjusting for sex and batch effects. Only 107 gene x cell type were significant after multiple-testing correction, of which 106 were in microglia cells (Fig. 3A; adj. *P* < 0.05 and |ρ| > 0.3).

**Fig. 3.**
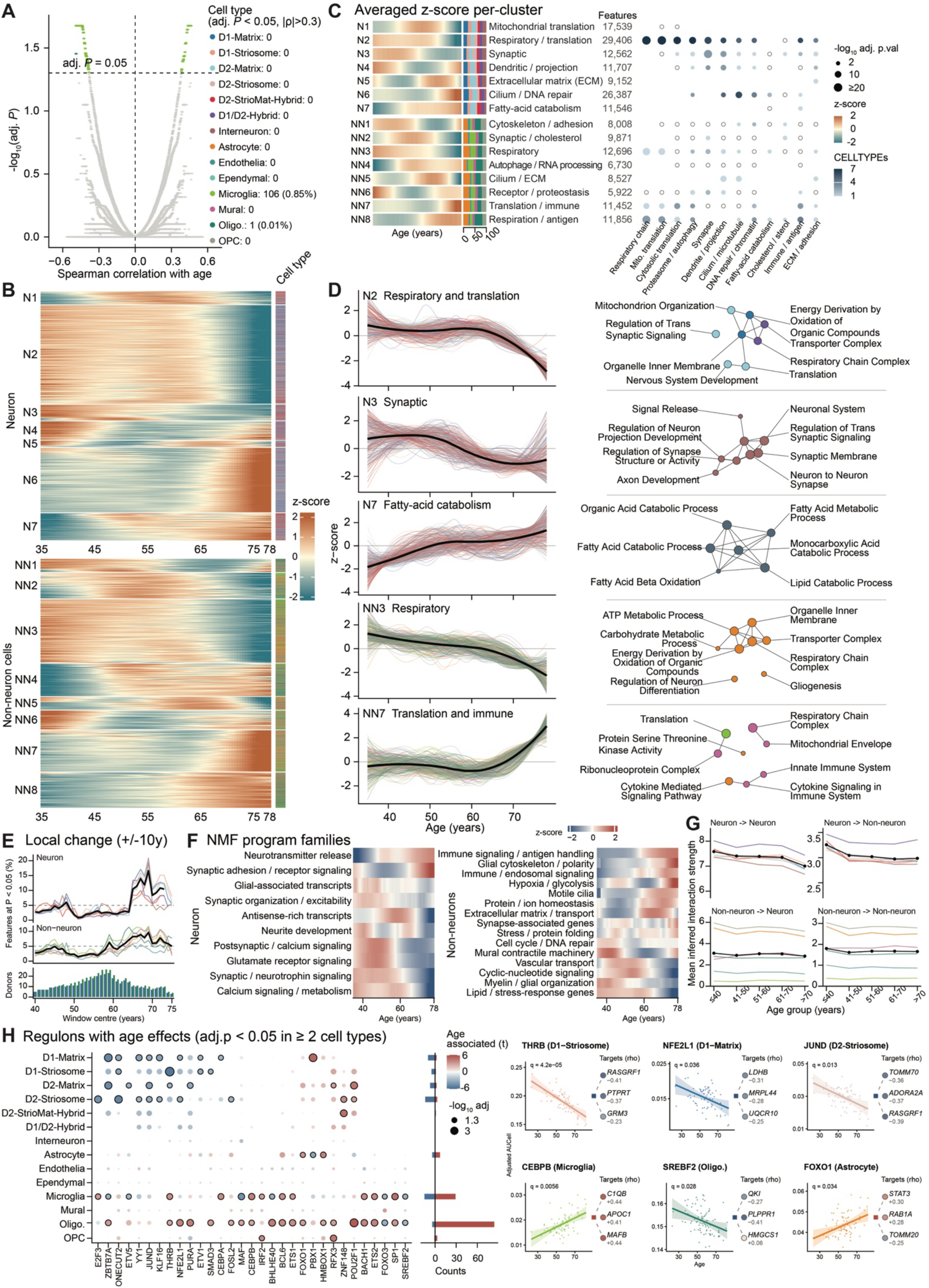
Molecular and cellular dynamics during healthy human caudate nucleus aging. **(A)** Age-expression linear correlations in 85 control donors. Points indicate genes with adj. *P* < 0.05, colors denote cell type. **(B)** Smoothed trajectories of gene x cell type features with age-group differences relative to donors aged ≤40 years (adj. *P* < 0.05). Rows are grouped by trajectory cluster; colors show *z*-scores, and side bars indicate cell type. **(C)** Summary of seven neuronal and eight non-neuronal trajectory clusters. Dot size indicates −log10(adj. *P*), and color indicates the number of cell types with significant enrichment. Open circles indicate whole-cluster enrichment only. **(D)** Selected representative trajectory clusters and pathway enrichment networks. Colored lines show individual feature trajectories; black lines show cluster centroids. Node color indicates cell type (gray, whole-cluster analysis), size indicates −log10(adj. *P*), and edges indicate gene-set overlap. **(E)** Local expression changes in ±10-year windows. Lines show the percentage of features with adj. *P* < 0.05 for individual cell types (colored) and pooled neuronal or non-neuronal features (black). The dashed line marks 5%. Bottom bars summarize donor counts. **(F)** Age trajectories of the MP families. Colors show the median standardized trajectory across MPs within each family. **(G)** CellChat-inferred interaction strength across age groups. Black lines show mean strength across cell type pairs; colored lines show means for each sender cell type. **(H)** Age associations of regulon activity with selected examples. Dot color shows moderated *t*-statistics; size is −log10(adj. *P*), and black outlines mark adj. *P* < 0.05. Bars show counts of increasing and decreasing regulons. Example plots show donor-mean AUCell activity with linear fits and 95% confidence intervals (CIs). Squares indicate regulon direction; circles show inferred targets, colored by their age-expression Spearman correlations.

To characterize the cell type-specific molecular dynamics of aging, we used smoothed expression trajectories^54^ to capture changes that emerged or accelerated during particular periods of adulthood (Fig. 3B). Clustering neuronal and non-neuronal gene x cell type features separately identified seven neuronal (N) and eight non-neuronal (NN) clusters. Some clusters changed gradually with age, while others remained relatively stable through midlife and then increased or decreased in later age windows (Fig. 3C), revealing substantial variation in the timing of age-associated expression changes.

Among the representative trajectories, the neuronal respiratory and translation cluster (N2) remained relatively stable through midlife and declined sharply at older ages (Fig. 3D). The NN3 cluster declined more gradually and was enriched for respiratory-chain genes. These patterns agree with reported decreases in respiratory and translation-related gene expression during human brain aging^55,56^. In contrast, N7 increased with age and was enriched for fatty acid catabolism and β-oxidation, extending the lipid-related expression changes observed in the previous section (Fig. 2E), consistent with recent reports of conserved lipid remodeling during aging^49^.

To locate the periods of greatest transcriptional change, we compared expression in sliding age windows (Fig. 3E). Neuronal changes showed a sharp increase in gene expression level around 65 years, with a peak in the late 60s, whereas non-neuronal changes increased more gradually (Fig. 3E). The atlas-defined MP families showed pronounced changes over a similar period, around 60-70 years, including declining neuronal synaptic and calcium-signaling programs and increasing non-neuronal immune programs (Fig. 3F). Inferred neuron-to-neuron interaction strength also showed an overall decline with age (Fig. 3G), consistent with the young-old comparison in Fig. 2F.

To identify regulatory programs associated with these expression changes, we examined age-dependent regulon activity (Fig. 3H). THRB regulon activity in MSNs and SREBF2 activity in oligodendrocytes declined with age, consistent with the shared young-old changes observed in Fig. 2H. NFE2L1, which supports neuronal proteasome function^57^, also showed declining regulon activity together with its inferred target genes *MRPL44* and *UQCR10*. These changes accompanied with the decline trend in respiratory and translation-related genes of cluster N2. Increased microglial C/EBPβ activity and C1QB expression paralleled the rise in immune programs, consistent with the role of C/EBPβ in inflammatory and complement-associated microglial responses^58^.

### Alcohol use disorder remodels caudate aging trajectories across molecular layers

To examine whether AUD alters normal caudate nucleus aging, we modeled donor-level pseudobulk expression in 81 controls and 82 individuals with AUD aged 24-82 years, including age, diagnosis (i.e., control or AUD), and their interaction, with adjustment for sex and batch effects (See Methods). Following the control-defined clusters from Fig. 3, we revealed widespread disruption of normal aging trajectories (Fig. 4A). The respiratory/translation cluster N2 showed a pronounced late-life decline in controls but remained relatively flat in AUD, while the late-life increase in fatty acid catabolism cluster N7 was replaced by a decline in older AUD donors. Individual genes also showed altered mitochondrial and lipid-related expression (Fig. 4B). *COX5B* and *UQCRQ*, encoding subunits of mitochondrial respiratory complexes IV and III, respectively^59^, decreased with age in control D2-striosome MSNs but increased in AUD. Lipid catabolism genes also decreased with age in AUD. These include *CPT1A* (supports mitochondrial fatty acid oxidation^60^) in D2-striosome MSNs, and *LIPA* (encodes the lysosomal acid lipase responsible for breaking down cholesteryl esters and triglycerides^61^) in OPCs.

**Fig. 4.**
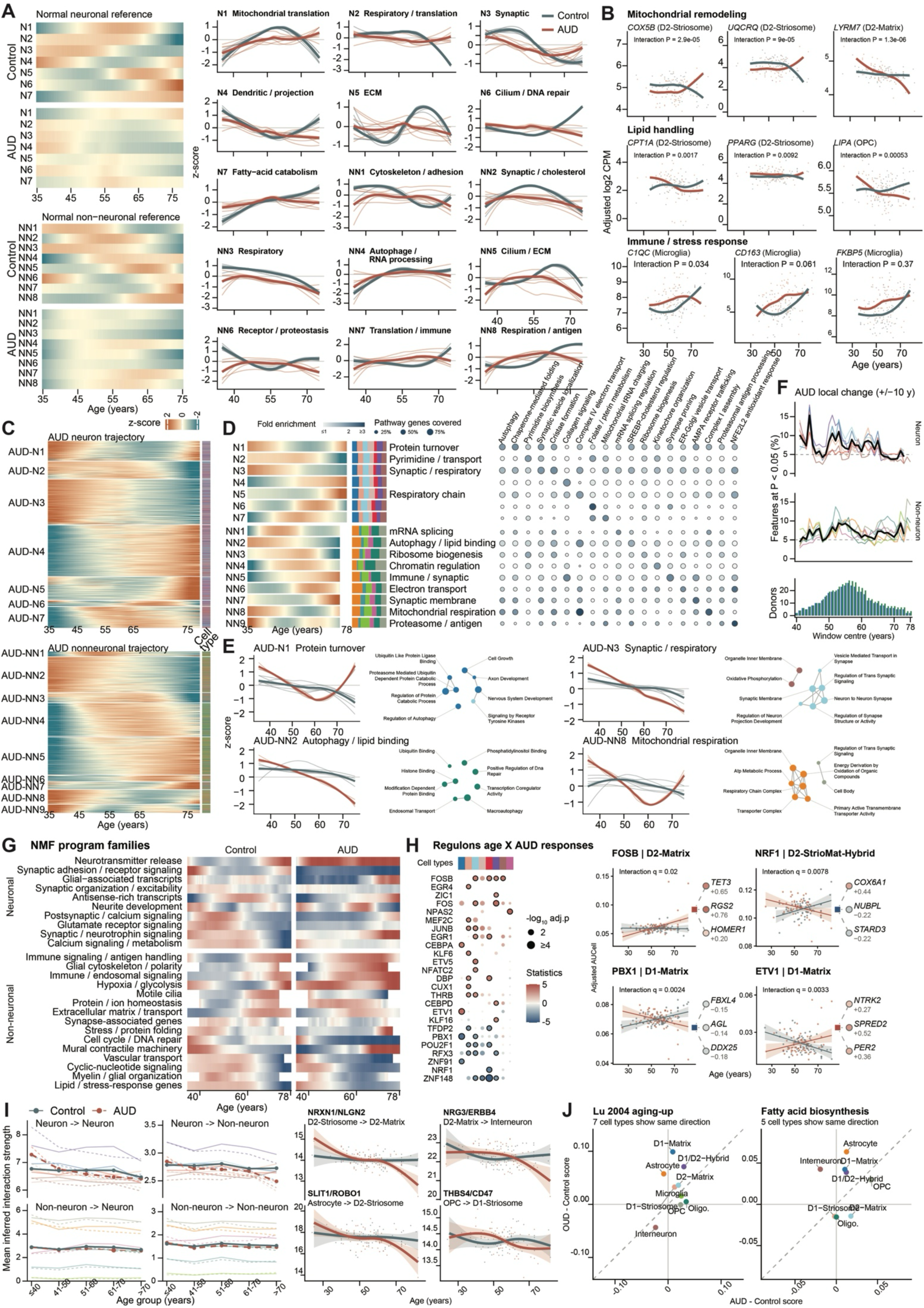
Alcohol use disorder remodels aging trajectories of the human caudate nucleus.(A) Control-defined aging clusters (as in Fig. 3B) in 81 control and 82 AUD donors. Left, heatmaps show mean cluster z-scores, standardized using control trajectories. Right, bold lines show cluster means; background lines show cell type means. Blue, control; red, AUD. **(B)** Selected gene-expression trajectories. Points represent donors; curves show smoothed fits for each diagnosis. Interaction *P* tests the difference in age slopes between controls and AUD. **(C)** Smoothed expression trajectories defined within AUD. Rows show gene x cell type features with age-group differences relative to donors aged ≤40 years (adj. *P* < 0.05), grouped by trajectory cluster. Colors show *z*-scores; side bars indicate cell type. **(D)** Summary of seven neuronal and nine non-neuronal AUD-defined clusters, showing cluster trajectories, cell type composition, and pathway enrichment. Dot color indicates fold enrichment; size indicates the percentage of pathway genes covered. **(E)** Selected AUD-defined trajectories with corresponding control curves, shown on the AUD scale. Lines are as in **(A)**. Network nodes represent enriched pathways; color indicates cell type, size indicates −log10(adj. *P*), and edges indicate gene-set overlap. **(F)** Local expression changes within AUD in ±10-year windows. Lines show the percentage of features with *P* < 0.05 for individual cell types (colored) and pooled neuronal or non-neuronal features (bold). Bottom bars summarize donor counts. **(G)** Age trajectories of the MP families in controls and AUD. Colors show median standardized MP scores within each family. **(H)** Age × AUD effects on regulon activity, with selected examples. Dot color shows moderated *t* statistics, size indicates - log10(adj. *P*), and black outlines mark adj. *P* < 0.05. Donor plots show AUCell activity with linear fits and 95% CIs. Squares indicate the direction of the regulon interaction; circles show corresponding effects on inferred target expression. **(I)** Inferred cell-cell communication. Left, CellChat interaction strengths across age groups; bold lines show means across cell type pairs, and thin lines show means for each sender cell type (solid, control; dashed, AUD). Right, selected donor L/R scores with fitted curves and 95% CIs. **(J)** Comparison of AUD and OUD effects on selected gene sets in donors aged 23-55 years. Points represent cell types; axes show age- and sex-adjusted disease-control score differences within each cohort. Dashed lines show equal values.

To identify the aging patterns emerging within AUD, we independently clustered AUD expression trajectories into seven neuronal and nine non-neuronal clusters with distinct functional enrichments (Fig. 4, C and D). The synaptic/respiratory cluster AUD-N3 declined more steeply than its matched control trajectory, encompassing synaptic genes across neuronal populations and oxidative phosphorylation genes in interneurons (Fig. 4E). The autophagy/lipid-binding cluster AUD-NN2 likewise showed a steeper decline across non-neuronal populations. These patterns suggest accelerated age-associated decline in selected synaptic, respiratory, and cellular maintenance programs^32^. Protein turnover cluster AUD-N1 and mitochondrial respiration cluster AUD-NN8 showed a different disruption: expression declined through midlife and rebounded at older ages, whereas the corresponding control trajectories declined gradually.

To determine when these changes occurred, we compared their distribution across adulthood. Neuronal transcriptional changes were greatest during midlife in AUD, particularly in the 40s and 50s (Fig. 4F), preceding the sharp increase around 65 years to the late 60s in controls (Fig. 3E). The atlas-defined MPs provided further evidence of earlier transitions (Fig. 4G). Immune/endosomal signaling shifted from lower scores (blue) to higher scores (red) during the 50s in AUD, compared with approximately the mid-60s in controls. Lipid/stress-response and myelin/glial organization programs showed earlier transitions in the opposite direction, from red to blue around 50-55 years in AUD and approximately 65 years or later in controls. Selected neuronal gene families, including synaptic/neurotrophin signaling, also reversed their normal age trends, increasing in AUD while declining in controls.

To identify regulatory changes accompanying these altered trajectories, we compared age-associated regulon activity in controls and AUD donors. The differences were concentrated in neurons (Fig. 4H). NRF1, a transcriptional activator of nuclear-encoded respiratory genes, showed increasing regulon activity in control D2-StrioMat-Hybrid MSNs and declining activity in AUD^62^. Its inferred targets NUBPL and STARD3 also had more negative age slopes in AUD. These genes encode a respiratory complex I assembly factor and an intracellular cholesterol transporter, respectively, connecting the regulatory changes to the respiratory and lipid-related trajectories above^63,64^.

To assess age-associated changes in potential cell-cell signaling, we compared inferred ligand-receptor interaction strength across age groups in control and AUD donors. Predicted cell communications, including neuron-to-neuron interactions, declined more strongly with age in AUD than controls (Figs. 3G and 4I). Donor-level scores decreased for NRXN1-NLGN2 between D2-Striosome and D2-Matrix MSNs^65^, and for NRG3-ERBB4 between D2-Matrix MSNs and interneurons^66^. These molecular pairs participate in synaptic adhesion and interneuron synapse organization^65,66^, suggesting disrupted regulation of neuronal connectivity during AUD aging. Declining SLIT1-ROBO1 and THBS4-CD47 scores further implicated communication from astrocytes and OPCs to MSNs.

Finally, to assess whether selected aging-related changes were shared with OUD, we compared both disorders with their respective controls among donors aged 23-55 years (Fig. 4J). The Lu2004 aging-up^6^ signature changed in the same direction in seven of nine cell types, including shared increases in four MSN populations, microglia, and oligodendrocytes. Fatty acid biosynthesis scores showed concordant changes in five of eight cell types. These findings identify shared aging-related expression changes in AUD and OUD, complementing previous evidence that the disorders converge on selected inflammatory and synaptic pathways^67^.

### Caudate nucleus aging programs recur across bulk brain cohorts and shift in disorder-specific directions

To assess shared and divergent aging signatures with other brain regions and disorders, we assembled 3,139 bulk RNA-seq tissue profiles from five cohorts^23,68–71^ covering 10 brain regions (Fig. 5A). These included BrainSeq Phase 3 (SZC and BIP)^23^, the Religious Orders Study and Rush Memory and Aging Project^69^ (ROSMAP), two AUD cohorts reported by Willis et al.^70^ and Lim et al.^68^, and an OUD cohort reported by Seney et al.^71^. The regions included the caudate nucleus, nucleus accumbens (NAcc), dorsolateral prefrontal cortex (DLPFC), and posterior cingulate cortex (PCC).

**Fig. 5.**
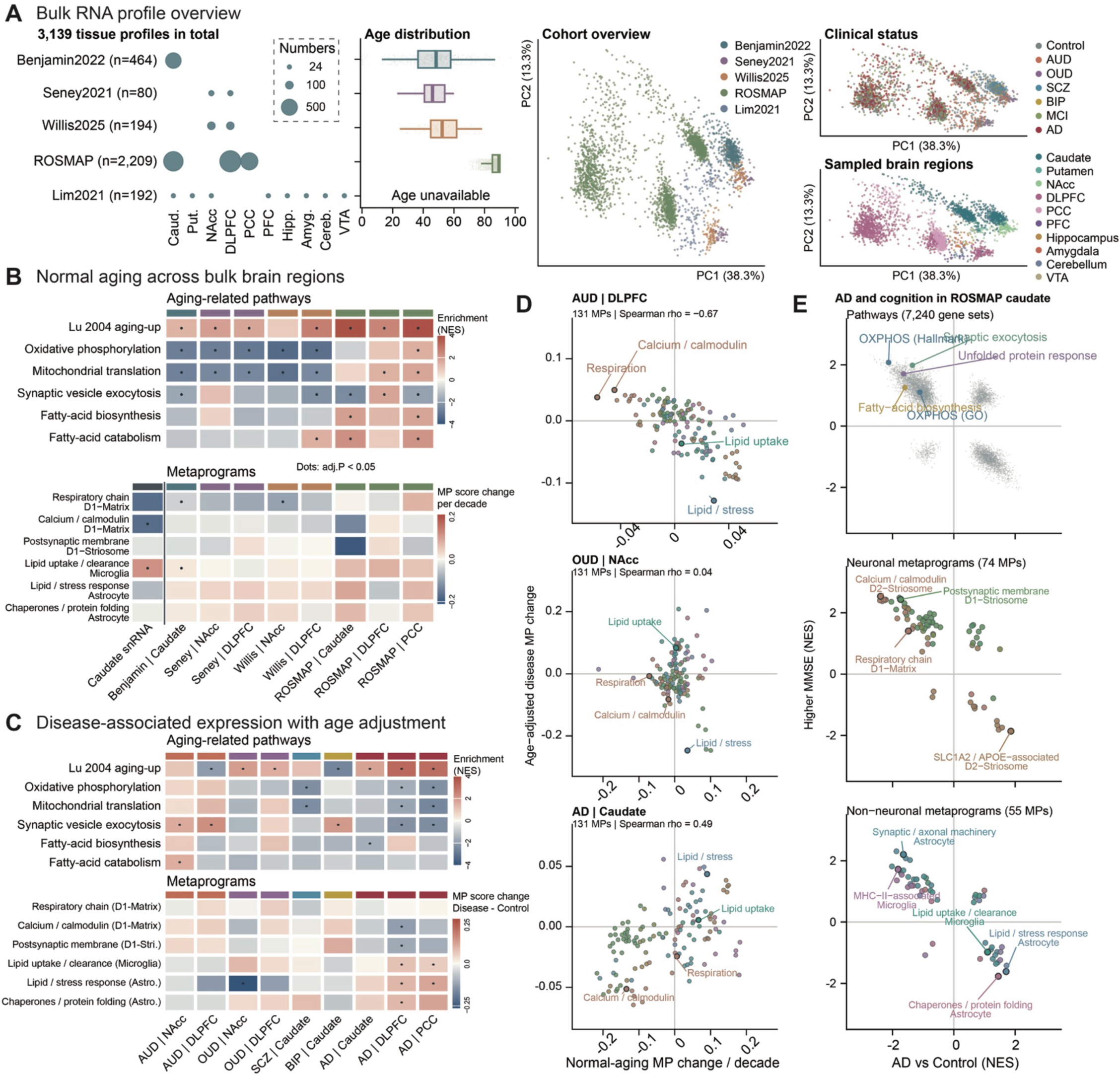
Caudate aging signatures across bulk brain regions, disorders, and cognition. **(A)** Overview of 3,139 bulk RNA-seq tissue profiles from five cohorts across 10 brain regions. Dot size indicates the number of profiles. Boxplots show age distributions. PCA plots are colored by cohort, clinical status, and brain region. **(B)** Normal-aging associations in controls. Top, normalized enrichment scores (NES) for selected aging-related gene sets. Bottom, MP score changes per decade; the first column shows the caudate snRNA-seq reference. Header colors indicate cohorts. Dots mark adj. *P* < 0.05. **(C)** Age-adjusted disease effects relative to controls from the same cohort and brain region. Colors show gene-set NES (top) and disease-control MP score differences (bottom). Rows correspond to **(B)**, and header colors indicate diagnoses. Dots mark adj. *P* < 0.05. **(D)** Comparison of normal-aging and disease effects across 131 human MPs. Each point represents an MP, plotted by its control age effect per decade (x-axis) and age-adjusted disease effect (y-axis). Colors indicate MP families; outlined points highlight selected examples. Spearman correlations are shown. **(E)** Associations with Alzheimer’s disease (AD) and cognition in ROSMAP caudate. Points show NES for AD versus control (x-axis) and higher Mini-Mental State Examination (MMSE) scores (y-axis). Top, pathways; middle and bottom, neuronal and non-neuronal MPs, colored by MP family. Selected gene sets are labeled. SCZ, schizophrenia; BIP, bipolar disorder; MCI, mild cognitive impairment. NAcc, nucleus accumbens; DLPFC, dorsolateral prefrontal cortex; PCC, posterior cingulate cortex; PFC, prefrontal cortex; VTA, ventral tegmental area.

We first asked which caudate aging features recurred across brain regions (Fig. 5B). The Lu2004^6^ aging-up signature showed positive enrichment across all eight cohort-region comparisons. Oxidative phosphorylation and mitochondrial translation decreased with age in the caudate, NAcc, and DLPFC in cohorts spanning broader adult age ranges, consistent with the respiratory and translation-related declines in caudate neurons (Fig. 3). The caudate-derived respiratory chain and calcium/calmodulin MPs also generally showed negative age associations, whereas the microglia-derived lipid uptake/clearance MP generally increased. Fatty acid biosynthesis and catabolism increased particularly in ROSMAP caudate and PCC samples. Together, these findings suggest that neuronal energy metabolism and signaling, together with glial lipid metabolism, change across brain regions during aging. This is consistent with previous report of reduced mitochondrial and synaptic gene expression in aging human cortex, and lipid-associated microglial dysfunction in the aging brain^6,49,55,72^.

We next asked whether disease-associated changes followed these aging patterns (Fig. 5C). The Lu2004^6^ aging-up signature showed positive enrichment in most disease comparisons, suggesting a broad overlap between disease-associated expression changes and normal brain aging. It also showed concordant changes in AD cortex: lower mitochondrial and synaptic signatures accompanied lower neuron-derived MP scores and higher glia-derived lipid MP scores. The recurrence of lipid-related programs in aging and AD points to glial lipid handling as a potential connection between them, supported by evidence linking lipid accumulation in AD microglia to neurotoxicity^73^. Interestingly, we found that aging-related mitochondrial function and fatty acid metabolism are consistently reduced in AUD, OUD, BIP, SCZ, and AD.

To assess how disease relates to normal aging, we compared disease and age effects across all 131 human MPs within the same cohort and region (Fig. 5D). These effects were inversely correlated in AUD DLPFC (Spearman ρ = −0.67), largely unrelated in OUD NAcc (ρ = 0.04), and positively correlated in the samples from the caudate nucleus from AD patients (ρ = 0.49). In AUD DLPFC, respiratory chain and calcium/calmodulin MP scores decreased with age but were higher in AUD, while the lipid/stress response MP increased with age but was lower in AUD. In AD caudate, calcium/calmodulin MPs decreased with both age and disease, whereas the lipid/stress response MP increased in both comparisons. These contrasting relationships suggest that disease-associated changes overlap with normal aging differently across disease-region contexts, paralleling the heterogeneity of brain aging observed across disorders and regions in neuroimaging studies^30^.

To examine the cognitive relevance of these molecular changes, we compared AD-associated enrichment with enrichment associated with higher last-available Mini-Mental State Examination (MMSE) scores^74^ in ROSMAP caudate nucleus samples (Fig. 5E). Across pathways and MPs, the two comparisons showed broadly opposing patterns. Hallmark oxidative phosphorylation, the caudate-derived respiratory chain MP, and calcium/calmodulin MPs from all six MSN populations were lower in AD and positively associated with MMSE. The astrocyte-derived lipid/stress response MP showed the reverse pattern, increasing in AD and associating with lower MMSE. These findings implicate mitochondrial and neuronal signaling programs with both AD diagnosis and cognitive performance. This is consistent with proteomic evidence that higher mitochondrial and synaptic protein abundance is associated with greater cognitive stability in older adults^75^.

### Chromatin accessibility trajectories of the caudate aging and their remodeling in AUD

To examine the chromatin changes during aging and in AUD, we analyzed paired single nucleus RNA-seq and assay for transposase-accessible chromatin using sequencing (ATAC)-seq profiles from the AUD cohort^28^. The atlas comprised 257,524 caudate nuclei from 161 donors, including 80 controls and 81 individuals with AUD (Fig. 6A). Three donors were excluded from age analyses, leaving 158 donors aged 24-82 years (see Methods).

**Fig. 6.**
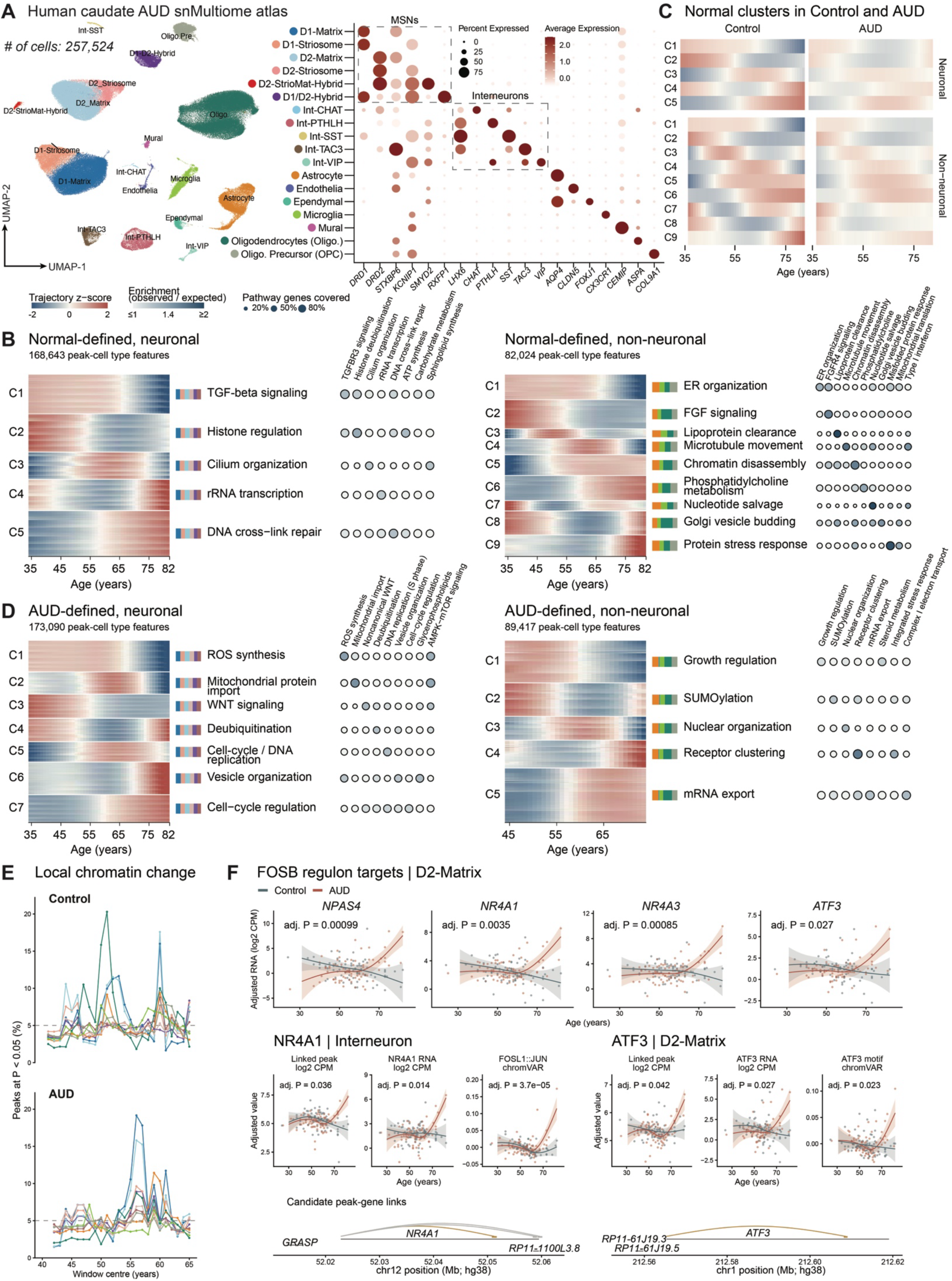
Age-associated chromatin responses in AUD connect to activity-dependent transcriptional programs. **(A)** Human caudate snMultiome atlas of 257,524 nuclei from 161 donors (80 controls and 81 AUD). UMAP colors indicate cell populations. Dot size indicates the percentage of nuclei expressing each marker, and color indicates average expression. **(B)** Control-defined chromatin accessibility trajectories and pathway enrichments. Heatmaps show trajectory z-scores for peak x cell type features grouped by cluster; bars indicate cell type contributions. Dot color indicates enrichment relative to matched background regions, and size indicates the percentage of pathway genes covered by core peaks. **(C)** Mean trajectories of control-defined clusters in controls and AUD, scaled to control donor means and SDs. **(D)** AUD-defined chromatin accessibility trajectories and pathway enrichments, displayed as in **(B)**. **(E)** Sliding-window changes in chromatin accessibility. Lines show the percentage of peaks differing between the older and younger halves of each age window (adj. *P* < 0.05), colored by cell type. **(F)** Age trajectories of selected FOSB regulon targets and linked chromatin features. Top, target-gene expression; bottom, linked-peak accessibility, gene expression, and chromVAR motif accessibility, with corresponding peak-gene links. Points represent adjusted donor values; lines and shaded bands show fitted trajectories and 95% confidence intervals. Slate and red indicate controls and AUD, respectively. Adj. *P* tests differences in age trajectories between groups.

We first defined chromatin accessibility trajectories during normal aging. Clustering control trajectories identified five neuronal and nine non-neuronal clusters, capturing gradual changes, midlife reversals, and pronounced late-life changes (Fig. 6B). A neuronal cluster associated with histone regulation and ATP synthesis declined with age, accompanying the respiratory and translation-related transcriptional declines in Fig. 3. Several clusters showed more pronounced changes around 65-70 years, including decreased accessibility in regions associated with TGF-β signaling and ER organization, and increased accessibility in regions associated with protein stress responses.

To determine how AUD alters these patterns, we followed the same control-defined clusters in AUD^28^. Their age-related changes were generally attenuated (Fig. 6C), paralleling the attenuation of normal transcriptional trajectories in Fig. 4A. Independent clustering of AUD trajectories revealed late-life accessibility gains in a neuronal cluster associated with vesicle organization and a non-neuronal cluster enriched for integrated stress-response genes (Fig. 6D). The neuronal vesicle-related cluster was also enriched for AP-1 motifs, connecting these accessibility changes to neuronal activity-related transcriptional responses^76^.

We next examined whether the timing of chromatin changes corresponded to the transcriptional transitions. In controls, local accessibility changes in D1- and D2-Matrix MSNs peaked around 60 years (Fig. 6E), close to the neuronal transcriptional transition during the 60s (Fig. 3E). In AUD, these accessibility peaks occurred in the mid-50s, paralleling the earlier transcriptional peak in Fig. 4F. Thus, neuronal chromatin remodeling and transcriptional changes showed a similar shift toward earlier adulthood in AUD.

To connect these accessibility changes to specific transcriptional programs, we examined paired RNA profiles and peak-gene links (Fig. 6F). In D2-Matrix MSNs, the expression of four inferred FOSB regulon targets, *NPAS4*, *NR4A1*, *NR4A3*, and *ATF3*, increased with age in donors with AUD. This pattern matched the age-related rise in FOSB regulon activity in the same cell type (Fig. 4H).

We next examined two individual loci: *NR4A1* in interneurons and *ATF3* in D2-Matris MSNs (Fig. 6F). At each locus, accessibility of the linked peak and expression of the corresponding gene increased in older AUD donors. The *NR4A1*-linked peak contained a FOSL1::JUN motif, and the *ATF3*-linked peak contained an ATF3 motif. Accessibility of regions containing these motifs also increased in older donors with AUD.

Together, the RNA and chromatin findings point to changes in activity- and stress-responsive gene regulation in older donors with AUD. The increased accessibility of regions containing the FOSL1::JUN motif is consistent with the role of AP-1 factors in activity-induced chromatin opening in response to neuronal activity^76^. Similarly, the up-regulation of *NPAS4* expression also suggests how neurons respond to activity: NPAS4 regulates synaptic gene programs involved in excitation-inhibition balance^77^.

## Discussion

We established cell type-resolved references for aging in the healthy human and macaque caudate nucleus and used them to examine their relationship to brain disorders, region, cognition, and chromatin regulation. Our findings show that caudate nucleus aging comprises genetic changes whose timing, direction, and conservation depend on cell type (Figs. 1-3). Across species, lipid-related gene sets and non-neuronal programs showed similar age-associated patterns, whereas neuronal programs often differed. The human neuronal programs showed more pronounced transitions in later adulthood than the more gradual changes in selected non-neuronal programs. AUD altered this organization by accelerating transitions, steepening declines in some programs, and attenuating or reversing others. Across additional brain cohorts, disease-associated expression changes showed convergent patterns, such as mitochondrial function and fatty acid metabolism, but largely different relationships to normal aging. Together, these observations support a view of caudate nucleus aging as a collection of partially shared cellular processes that can be altered differently by disease or behaviors.

The human-macaque comparison revealed both shared and species-dependent patterns of caudate aging. Age-associated metaprogram changes were positively correlated between species in microglia, oligodendrocytes, and endothelial cells, supporting shared features of non-neuronal aging (Fig. 2). Neuronal expression changes were more divergent, although inferred neuron-to-neuron communication declined in both species and THRB regulon activity showed a shared downward trend in MSNs. These findings indicate that agreement at the level of expression programs and inferred signaling can coexist with limited overlap among individual differentially expressed genes. They also suggest that the relevance of macaque models to human caudate aging should be evaluated for specific cell types and molecular processes. However, the small, female-only macaque cohort and differences in the ages and lifespan stages sampled limit the interpretation of cross-species differences. Broader sampling will be needed to determine whether the divergent neuronal patterns reflect species-specific aging or differences in the timing of comparable changes.

Human caudate aging showed distinct temporal patterns across cell types and molecular programs. A neuronal gene cluster enriched for respiratory-chain and translation functions remained relatively stable through midlife before declining in later adulthood. Neuronal synaptic and calcium-signaling programs also declined in later life, whereas microglial immune programs increased more gradually (Fig. 3). These patterns indicate that age-associated molecular differences vary across adulthood and may be incompletely captured by a single linear age association. The concentration of neuronal changes in later adulthood suggests more pronounced transcriptional remodeling during this period, particularly in programs related to energy metabolism, protein synthesis, and signaling. However, we inferred trajectories from cross-sectional comparisons, and their estimated timing depends on age coverage, donor variability, and the analytical approach. They therefore identify periods for closer investigation rather than establish a fixed transition age or a common trajectory followed by every individual.

AUD was associated with a reorganization of these age-dependent patterns (Figs. 4). Earlier transcriptional changes and earlier accessibility transitions in matrix MSNs are compatible with the acceleration of brain aging reported in cases of alcohol dependence^32^. Nevertheless, a uniform acceleration model does not explain the full set of findings. Some synaptic, respiratory and cellular maintenance programs declined more steeply in AUD, whereas other control-defined trajectories became flatter or reversed direction. In particular, the attenuation of a normal late-life decline cannot by itself be interpreted as preservation of function: it may reflect a different baseline, altered regulation, or differences in the individuals represented at older ages. The combination of these patterns indicates that AUD changes the coordination of molecular aging across programs, rather than simply moving the caudate along a single normal-aging trajectory more rapidly.

Changes in mitochondrial and lipid-related programs provide a biological connection between the altered trajectories and the broader disease comparisons (Fig. 5). Respiratory-chain and calcium/calmodulin programs generally declined with age across adult-spanning cohorts, while lipid-related programs showed cell type- and context-dependent changes. Their recurrence across disorders suggests that energy metabolism and lipid handling are common components of the molecular response to aging and disease. However, recurrence of a pathway does not imply a common direction of change or a shared causal mechanism. Across metaprograms, disease effects aligned with normal aging in the caudate nucleus of the AD cohort, opposed aging in AUD prefrontal cortex and showed little correspondence with aging in OUD NAcc. These differences argue against treating all disease-associated molecular changes as evidence of accelerated aging and emphasize the importance of the brain region and cellular program examined.

The associations with cognition extend the relevance of these molecular findings to cognitive performance (Fig. 5). In ROSMAP caudate, oxidative phosphorylation, respiratory-chain, and MSN-derived calcium/calmodulin signaling programs were lower in AD cases than in cognitively unimpaired controls, and positively associated with MMSE scores across donors, spanning diagnostic groups. The astrocyte-derived lipid/stress-response program showed the opposite pattern, with higher enrichment in AD and an association with lower MMSE scores. These findings link disease-associated differences in mitochondrial and neuronal signaling programs to cognition, consistent with evidence associating higher mitochondrial and synaptic protein abundance with greater cognitive stability in older adults^75^. However, they do not establish that the molecular differences cause cognitive impairment or that their relationships with cognition are specific to AD. Because these analyses used bulk tissue, differences in cell composition may also contribute to the observed associations. Cell type-resolved studies linked to longitudinal cognitive assessments will help determine which programs track cognitive decline and whether their associations persist after accounting for cellular composition. Paired RNA and chromatin profiles further suggest that altered neuronal aging in AUD involves activity- and stress-responsive regulation (Fig. 6). Coordinated increases in linked-peak accessibility and expression at NR4A1 and ATF3, together with AP-1-associated accessibility changes, identify candidate regulatory relationships for experimental testing. These observations are consistent with known roles of AP-1 in activity-induced chromatin opening^76^. However, motif enrichment and peak-gene correlations do not demonstrate transcription factor binding or establish the direction of regulation. The observed responses could reflect adaptation to chronic exposure, withdrawal-related activity, or cellular stress. Likewise, expression-based estimates of cell communication identify changes in signaling-related transcripts rather than direct changes in synaptic connectivity or transmission. Perturbation studies combining regulatory measurements with neuronal physiology will be necessary to establish their functional consequences.

Our analyses did not fully account for co-occurring disorders and mixed neuropathologies. AUD and OUD may co-occur with each other^78^ or with psychiatric disorders^79^. Such overlap could contribute to molecular signatures attributed to the primary diagnosis and complicate comparisons with normal aging. More detailed clinical histories and neuropathological assessments will be needed to distinguish disorder-specific effects from those associated with co-occurring conditions.

## Materials and Methods

### Human donors

Human caudate nucleus snRNA-seq data came from two published postmortem cohorts. The AUD cohort comprised 168 donors (84 with AUD, 25-82 years; and 84 controls, 24-82 years) profiled in 17 genotype-multiplexed pools^28^. The OUD cohort comprised 10 donors: 6 with OUD (35 to 54 years; 3 female, 3 male) and 4 controls (23, 39, 54 and 55 years; 3 female, 1 male)^29^. Of the 178 donors (42 female, 136 male), 173 had a recorded age (23 to 82 years). The five donors without age (139 nuclei) were kept in the atlas and composition summaries but excluded from all age analyses.

### Rhesus macaque samples

We collected caudate tissue from six female rhesus macaques (*Macaca mulatta*), including three young (5.0, 5.75 and 6.5 years) and three old (18.0, 19.0 and 19.5 years) from the Emory National Biomedical Research Center. All animal procedures followed the National Institutes of Health Guide for the Care and Use of Laboratory Animals and were approved by the Emory University Institutional Animal Care and Use Committee. Animals were anesthetized with ketamine/xylazine, sacrificed with an overdose of pentobarbital (100 mg/kg, i.v.), followed by removal of the brain. We blocked the brain with a matrix and isolated the caudate nucleus.

We generated a single nucleus suspension, which was then run through a 10X chromatin controller, followed by the generation of libraries (with 10X Genomics, v4 chemistry, Product Code 1000691), as previous published^12,80^. We sequenced the libraries at a depth of 150,000 per cell at the Genomics Core at the Emory National Biomedical Research Center.

### snRNA-seq pre-processing, integration, and annotation

Human UMI matrices were taken as processed by the source studies^28,29^. Macaque reads were aligned to rheMac10 with Cell Ranger (v10.0.0), and ambient RNA was removed with SoupX^81^. Genes were matched across species by symbol. Nuclei that had ≥1,200 detected genes and less than 10% mitochondrial reads were retained. The human and macaque atlas was integrated separately using Seurat v5^82^ package. We annotated the clusters according to published cell type-specific markers^12,80^.

### Pseudobulk profiles and statistical framework

For each donor and cell type, we summed the expression counts for each gene across all nuclei of that cell type. We included profiles with ≥15 nuclei, generating 2,038 human profiles from 175 donors and 84 macaque profiles from all six animals. For each analysis, genes were filtered with edgeR filterByExpr^83^, counts were TMM-normalized^84^, and normalized to log2 counts per million (log-CPM). Platform-origin batches were included as a covariate (17 AUD-cohort batches^28^ and the OUD cohort^29^).

In continuous models, age was expressed in decades, For age-group comparison, donors were grouped as ≤40, 41 to 50, 51 to 60, 61 to 70 and >70 years. For each sliding window, we compared donors in the 10 years below the window centered at age c with donors from the center through 10 years above it, namely [c - 10, c) vs. [c, c + 10]. Donor-level scores were analyzed with limma^85^ and robust empirical Bayes moderation^86^. *P* values were adjusted by the Benjamini-Hochberg (BH) procedure^87^ within each species × cell type and contrast.

### NMF metaprograms

Metaprograms (MPs) were derived as described^38^. Consensus nonnegative matrix factorization (cNMF)^39^ was run on each donor × cell type with ≥30 nuclei (1,804 human and 81 macaque runs), after removal of mitochondrially encoded and ribosomal protein genes, over the 3,000 most overdispersed of the 5,000 most highly expressed genes (K = 4 to 9). Each program was represented by its 50 top-loading genes and retained when it recurred within the donor across K (≥30 shared genes) and in another donor (≥10 shared genes). Within each species and cell type, robust programs were clustered greedily at ≥10 shared genes into MPs of the 50 most frequent genes. MPs were annotated by over-representation against MSigDB Hallmark, GO and C8 sets^88^ and SynGO^89^ and labeled by manual review. MP families came from average-linkage clustering of 1 - Jaccard similarity across both species, separately for neuronal and non-neuronal MPs. Within a cell type, a human and a macaque MP were matched when they shared ≥10 of their 50 genes (hypergeometric test; BH across 701 pairs). Donor MP scores used the control-gene method^90^ on gene-centered log-CPM. Each MP was scored in its discovery cell type, and all 199 MPs were also scored in all 14 cell types of both species.

### Gene regulatory networks

Regulons were inferred separately for each species with pySCENIC^45,46^ (v0.12.1), using nuclei of all donors and diagnoses. An atlas catalogue was built from per-donor GRNBoost2^91^ networks (donors with ≥50 nuclei), and per-cell type catalogues from donor-balanced references of about 5,000 nuclei. Activating modules were pruned with cisTarget against hg38 v10 motif rankings (normalized enrichment score ≥3). AUCell^45^ scored unweighted regulons in every nucleus, and scores were averaged per donor and cell type. Atlas regulon specificity was measured by the regulon specificity score^92^. Regulons were matched between species by transcription factor (TF) and direction; target overlap was tested in a common universe excluding the TF (one-sided hypergeometric test), and activity profiles were compared by Spearman correlation of cell type means.

### Young-vs-old comparison

The cross-species design followed Jostard *et al.*’s study^93^. To estimate the human-equivalent ages of the macaque groups, we compared pooled macaque profiles with human control profiles over the 3,000 most age-correlated genes. The estimations were calculated per cell type, using human age range of 15-35 years for the young macaque group and 55-85 years for the old macaques. These estimates guided the selection of human controls for the young-vs.-old comparison: young human group, aged ≤37 years (n = 6); aged human group, 65-78 years (n = 13). Effects were reported as old minus young. Cell type proportions were compared between age groups after adjusting for sex in humans.

Differential expression used one joint limma-voom model per species^85,94^, ∼ 0 + group:celltype:sex + n_cells + gdr (number of nuclei and standardized gene detection rate), on genes with mean count >5 (18,693 human and 19,179 macaque genes). Sample quality weights^95^, within-donor correlation^96^, and surrogate variables^97^ were estimated before the final fit. Cell type contrasts averaged the sex strata, and q values were computed with swfdr^98^ jointly across the 14 cell type contrasts and three composites (all, neuronal and non-neuronal). MP scores in all 28 species × cell type contexts and donor-mean AUCell scores were modeled as ∼ group + sex (human) or ∼ group (macaque) in contexts with ≥3 donors per group (BH across MPs or regulons within context), and human and macaque MP changes were compared by Spearman correlation.

### Normal aging in controls

Aging dynamics were analyzed following Shen *et al.*’s design^54^ in the 85 controls with recorded age (23 to 82 years). Within each cell type, log-CPM was adjusted for sex and pool by linear regression, and the residuals were correlated with age (Spearman; BH adjustment within cell type). Each older age group was compared with the ≤40 group by Wilcoxon tests (≥3 donors per group), and features with adj. *P* < 0.05 in any comparison were called stage-changed.

For the sliding-window analysis, a modified DE-SWAN^54,99^ compared younger and older donors within overlapping ±10-year windows. Windows centered ranged from 40 to 75 years in one-year steps (Wilcoxon test; ≥3 donors per side), and local change was the fraction of features at adj. *P* < 0.05 per cell type and compartment. Trajectories were LOESS fits of adjusted expression on age (span 0.9, chosen by cross-validation; 35 to 78 years), z-scored and clustered jointly within each compartment (118,299 neuronal and 75,062 non-neuronal gene × cell type features; ependymal cells excluded) by fuzzy c-means^100^, with the number of clusters at the elbow of the minimum centroid distance^101^. Features with membership >0.5 formed the cluster cores. The same procedure applied to each cell type separately gave 85 clusters, used only as bulk scoring references.

MP scores (discovery cell type) and donor-mean AUCell^45^ scores were modeled as ∼ age10 + sex + pool (BH within cell type). MP trajectories were LOESS fits of scores adjusted for sex and pool, scaled by the mean and SD of the fitted curve; family curves were the median across member MPs, and family change was the difference in mean z between ≥70 and ≤45 years. Regulon targets that changed in the regulon’s direction (Spearman adj. *P* < 0.05) were tested for over-representation among the tested targets of all regulons of the cell type (hypergeometric test; BH adjustment^87^ across 192 regulons × cell type pairs).

### Age × AUD interaction

AUD-cohort donors with recorded age (81 controls and 82 with AUD; 24 to 82 years) were modeled per cell type with voom-limma^85,94^ as ∼ age*AUD + sex + pool. Four contrasts were estimated: the control and AUD age slopes, their difference (the age × AUD interaction) and the AUD - control difference at 50 years. Each contrast was also tested with CAMERA^102^ (GO, KEGG and Reactome sets; BH adjustment^87^ within cell type × contrast).

Within each diagnosis, log-CPM adjusted for sex and pool was analyzed with the Spearman, age-group and window tests of the control analysis and fitted by LOESS over ages shared by both diagnoses (35 to 78 years). Fig. 3 cluster memberships were held fixed, and both diagnoses were scaled by the mean and SD of each feature’s control curve. Within each cell type, normal cluster and sign, core genes with slope-difference adj. *P* < 0.05 were tested for over-representation against the tested genes of all normal clusters of the cell type (hypergeometric test; BH adjustment within cell type × cluster × sign). AUD-defined clusters were derived from AUD curves scaled by their own mean and SD, with matched control curves on the same scale. MP scores (discovery cell type) and AUCell scores were modeled with the age × AUD design (BH adjustment within cell type × contrast); MP trajectories with both diagnoses scaled by the control curve; and regulon targets with slope differences of the regulon’s sign (adj. *P* < 0.05) were tested as above (hypergeometric test).

To compare AUD with OUD, two gene sets were scored with the control-gene method: the brain aging-up genes (264 genes)^6^ and GO fatty acid biosynthetic process (180 genes). Normalization, gene centering and control genes used all profiles of the cell type across cohorts, diagnoses and ages, and a set required ≥10 scored genes covering ≥50% of its members. In donors aged 23 to 55 years, each disorder was compared with the controls of its own cohort as score ∼ diagnosis + age (per decade) + sex, plus pool in the AUD cohort (robust moderation), in cell types with ≥3 donors per group in both cohorts.

### Functional enrichment

All enrichment analyses used one gene-set catalogue from MSigDB^88^ (2025.1.Hs) and SynGO^89^ (v2023-12-01). Catalogue symbols absent from the data were mapped to data symbols through one- to-one NCBI Gene aliases in the snRNA-seq analyses of Figs. 2 to 4 and through unambiguous aliases in bulk data. Ranked enrichment^103^ used fgseaMultilevel on moderated t statistics with Hallmark, GO, Reactome, KEGG, WikiPathways and SynGO (15 to 500 tested genes; BH adjustment within ranking). Over-representation analysis (fgsea fora) tested gene lists against their tested universes with GO, KEGG and Reactome sets of 15 to 500 genes (BH within list). Whole-cluster tests pooled the unique core genes of a trajectory cluster across cell types and tested them against all clustered genes of the compartment. The brain aging-up and aging-down gene sets^6^ and the seven top-level MitoCarta3.0^104^ were tested separately. For Fig. 2C, GO biological process, Reactome and KEGG legacy pathways with adj. *P* < 0.05 in any cell type of either species were linked at Jaccard index ≥0.10. They were grouped by weighted Louvain clustering (igraph; seed 1), and communities of ≥10 pathways were labeled. Trajectory-cluster terms at adj. *P* < 0.05 were assigned to function themes by keyword matching of term names. Cluster networks showed up to eight terms linked at Jaccard ≥0.1.

### Cell-cell communication

CellChat^105,106^ (v2.2.0) was run on log-normalized expression using the CellChatDB.human database. Networks were built for young and old humans and macaques (Fig. 2), the five control age groups (≤40, 41-50, 51-60, 61-70, and >70 years; Fig. 3), and each diagnosis × age group of the AUD cohort (Fig. 4). Within each figure, every cell type contributed the same number of nuclei to each network, drawn in rotation across donors, and communication probabilities used the trimean with 100 permutations (*P* < 0.05). Summaries were outgoing and incoming strength, Old - Young changes in edge weight and in summed neuron-to-neuron probability for the 147 ligand-receptor (L/R) pairs retained among neurons in both species (Fig. 2), and mean summed probability per directed cell type pair within neuronal and non-neuronal blocks (Figs. 3 and 4).

For Figs. 2 and 4, each sender-receiver-L/R combination significant in any network was scored per donor as the mean log-CPM of the ligand subunits in the sender plus that of the receptor subunits in the receiver (≥15 nuclei each). Scores were modeled with the young-versus-old designs (BH within species) or per sender-receiver pair with the age × AUD design (BH within contrast), and displayed as natural-spline fits with pointwise 95% CIs after removal of sex and batch effects.

### Bulk RNA-seq cohorts

We reanalyzed gene-level counts of 3,139 postmortem profiles from five cohorts^23,68–71^: BrainSeq Phase 3 caudate with schizophrenia and bipolar disorder (Benjamin2022^23^); ROSMAP caudate, dorsolateral prefrontal cortex (DLPFC) and posterior cingulate cortex (PCC) with the final consensus cognitive diagnosis^69^; nucleus accumbens (NAcc) and DLPFC with AUD (Willis2025^70^) and with OUD (Seney2021^71^); and eight regions with AUD and no individual ages (Lim2021^68^). Within each of 16 cohort × region blocks, genes were filtered and TMM-normalized, and principal component analysis (PCA) used the 5,000 most variable of the 12,667 genes retained in all blocks, without cross-cohort correction. The association of PC1 with region, cohort and clinical status was summarized by R^2^.

Models used adults (≥18 years), one profile per donor and region, and each cohort’s own controls, with age per decade centered at the model mean (age10). Control-aging models were ∼ age10 + covariates in controls with exact ages (eight contexts; ROSMAP donors aged ≥90 years excluded). Disease models were additive, ∼ group + age10 + covariates; in ROSMAP (AD, cognitive diagnosis 4 versus 1), age was capped at 90 with a ≥90 indicator; Lim2021 had no age information thus was excluded. Covariates were sex and postmortem interval (PMI), RNA integrity number (RIN), batch and race where recorded and variable, and depression and smoking in Willis. Genes were fitted with voom^94^ and robust moderation^86^ (BH within model and contrast), and ranked enrichment used each model’s moderated t statistics. Reference programs were scored in each profile with the control-gene method and tested with the same designs: the 131 MPs, the 85 per-cell type trajectory-cluster cores, two sets of microglial genes associated with age in controls (rising or declining) and up to nine cell type marker sets (≤50 genes per cell type by the marker rule above), with BH across the 226 or 227 programs per model. Control-aging and disease effects of the 131 MPs in the same cohort and region were compared by Spearman correlation.

For ROSMAP caudate (Fig. 5E), AD (n = 250) was compared with no cognitive impairment (n = 248) as ∼ diagnosis + ns(age, 3) + ≥90 indicator + sex + PMI + race, and last-visit Mini-Mental State Examination score^74^ (MMSE) (per 5 points) was modeled in 743 donors of all diagnoses with the same age terms plus sex, education, PMI, race and study (ROS or MAP). Genes ranked by moderated t were tested against the catalogue (BH within ranking) and for the 129 MPs with ≥15 measured genes (BH across MPs).

### snMultiome data

Paired snRNA-seq and snATAC-seq profiles (10x Genomics Multiome) of AUD-cohort caudate nuclei^28^ were analyzed from the processed object (257,524 nuclei; 30,862 common peaks; hg38), with interneuron subtypes merged. Of the 161 donors, two were excluded because age was unavailable, and one donor labeled AUD/WE was excluded because the analysis was restricted to Control and AUD groups, leaving 158 donors. RNA and peak counts were summed per donor and cell type; profiles with ≥15 paired nuclei came from 158 donors (79 controls and 79 with AUD; 24 to 82 years). A cell type was analyzed if had ≥20 donors per diagnosis and distinct ages spanning ≥20 years. Sex and batch effects were considered. chromVAR^107^ deviations of 879 JASPAR 2024^108^ vertebrate motifs were computed per cell type from donor pseudobulk peak counts. RNA and peak counts were filtered, TMM-normalized and voom-weighted^94,95^; motif deviations had unit weights.

Age was modeled as a 3-df natural spline (knots at control-age tertiles), reparameterized as a linear term, age10, and two orthogonal nonlinear terms. Control models were feature ∼ sex + pool + spline(age); joint models added diagnosis and diagnosis-specific spline terms. After robust moderation, nested F-tests assessed age (3 df) and nonlinearity (2 df) within each diagnosis and the AUD - control curve difference (3 df) (BH within cell type × modality × model × test). Curves were evaluated at ages with ≥5 donors of a diagnosis within ±10 years, with pointwise 95% CIs. Within each diagnosis, peaks were also tested for linear age associations (∼ sex + pool + age10) and for local change between the older and younger halves of ±10-year windows (centers 40 to 65 years; ≥5 donors per side; BH within cell type × diagnosis × center).

Control-model curves formed the normal atlas and joint-model AUD curves the AUD atlas; curves were standardized and clustered per compartment by fuzzy c-means as above. With normal-atlas memberships fixed, control and AUD curves were scaled by the mean and SD of adjusted control values and compared by Spearman correlation. For pathway annotation, core regions were assigned to genes by promoter overlap (TSS -2 kb to +500 bp) or the nearest TSS within 250 kb, and the deduplicated core regions of each cluster, pooled across cell types, were compared with 1,999 permutations of atlas regions that preserved region counts within strata of GC content, accessibility, length, gene-assignment opportunity and assignment multiplicity. GO biological process, Reactome and KEGG sets of 5 to 500 assigned genes were tested (empirical one-sided P; BH within cluster; enrichment = (observed + 0.5)/(expected + 0.5)). One label per cluster was chosen among pathways with ≥10 genes, enrichment >1 and permutation P ≤ 0.01; all had adj. *P* > 0.05 and are descriptive. Core peaks were tested for motif enrichment against GC-, length- and accessibility-matched background peaks (hypergeometric test).

Signac^109^ LinkPeaks correlated accessibility with expression in up to 50 paired nuclei per donor for peaks within 250 kb of each TSS; links with adj. *P* < 0.05 and |r| > 0.05 were retained (2,606 links). Features with q < 0.05 for control age, AUD age or curve difference were grouped by k-means within each cell type and modality, and RNA groups were tested for over-representation of per-cell type regulon targets (hypergeometric test; BH across regulons). Fos- and Jun-related motifs were the 46 JASPAR matrices with these family annotations. Example loci were chosen after testing among links with curve differences at adj. *P* < 0.05 for both peak and gene and a same-sign donor-level correlation, taking the two genes with the largest smaller value of the peak and RNA curve ranges (NR4A1 in interneurons and ATF3 in D2-Matrix MSNs); each displayed motif had a sequence match in the linked peak. The frequency of immediate early genes among the 15 links on regions with AUD age or curve-difference effects was compared with that among all 2,606 links (hypergeometric test).

### Software

Analyses used R 4.5.1 (limma 3.64.3, edgeR 4.6.3, fgsea 1.34.2, msigdbr 25.1.1, sva, swfdr, e1071 1.7-17, irlba 2.3.5.1, igraph 2.1.4, CellChat 2.2.0, presto 1.0.0, AUCell 1.30.1, Seurat 5.3.0, Signac 1.16.0, chromVAR 1.30.1, motifmatchr 1.30.0 and JASPAR2024 0.99.7). Python 3.10 environments ran pySCENIC 0.12.1, arboreto 0.1.6 and cNMF 1.7.

## Data sources, code availability

Human AUD snRNA-seq and snMultiome data were obtained from GSE277313^28^. Human OUD snRNA-seq data are available at GSE225158^29^. Bulk data are available from BrainSeq Phase 3^23^, ROSMAP (AD Knowledge Portal^69^), and GEO (GSE253155^70^, GSE174409^71^, and GSE181982^68^). We will deposit macaque snRNA-seq processed and raw data to GEO. Code is available at https://github.com/JHELab2024/caudate_aging_and_disease.

## Acknowledgement

This research is supported by the Emory National Biomedical Research Center Grant No. ORIP/OD P51OD011132.

